# Regulatory effects on virulence and phage susceptibility revealed by *sdiA* mutation in *Klebsiella pneumoniae*

**DOI:** 10.1101/2024.09.27.615489

**Authors:** Sergio Silva-Bea, Pablo Maseda, Ana Otero, Manuel Romero

## Abstract

The World Health Organization has identified multi-drug resistant (MDR) *Klebsiella pneumoniae* strains as the highest priority in 2024. SdiA, a LuxR-like quorum sensing (QS) receptor that responds to *N-*acyl-homoserine lactones (AHLs), exerts a substantial regulatory influence on the virulence of numerous Gram-negative bacteria. The function of this receptor in the virulence of *K. pneumoniae* remains uncertain. Nevertheless, further investigation into the significance of this receptor is needed, as it represents an intriguing avenue with the potential to contribute to the development of novel antimicrobial strategies. The objective of the present study was to elucidate the function of SdiA in *K. pneumoniae* biofilm formation and virulence. To this end, a genetic knockout of *sdiA* was conducted, and virulence-related phenotypic studies were performed following AHL provision. The results demonstrate that SdiA deficiency increases susceptibility to phage infection and human serum resistance, and promotes biofilm maturation and cell filamentation. No effect on virulence was observed in vivo in the *Galleria mellonella* infection model. The addition of *N*-hexanoyl-L-homoserine lactone (C6-HSL) promoted SdiA-dependent biofilm maturation but also enhanced serum resistance and reduced virulence against *G. mellonella* in the absence of SdiA. The results of this study demonstrate that C6-HSL and SdiA exert a dual influence on virulence phenotypes, operating both independently and hierarchically. These findings provide new insights into the virulence of *K. pneumoniae* and its regulation by SdiA.

**Importance:** This study represents a significant contribution to our understanding of the complex regulatory mechanisms that govern the virulence of multi-drug resistant *Klebsiella pneumoniae* through quorum sensing (QS). The study offers insights into the function of SdiA, a QS receptor, in the regulation of biofilm formation, susceptibility to phage infection, serum resistance, and cell filamentation in this bacterium. Furthermore, the findings of this study demonstrate that exogenous *N*-acyl-homoserine lactone (AHL) signalling influences the aforementioned virulence phenotypes in both a SdiA-dependent and independent manner, as well as in a hierarchical manner.

## 3 Introduction

The emergence of MDR bacteria represents a significant public health concern, given the lack of effective treatment options (1). *K. pneumoniae* has been classified by the WHO as a maximum priority pathogen for the development of new antimicrobial strategies in 2024, with an increasing incidence of convergent strains with multirresistance and hypervirulence traits (2,3).

A comprehensive understanding of the factors that regulate virulence traits is essential for the development of novel antimicrobial therapies. Among these, antimicrobial therapies are being explored based on the hypothesis that blocking QS systems may play an important role in controlling virulence in many MDR pathogens. QS systems are responsible for regulating gene expression in bacterial populations in accordance with cell density through the production of autoinducing molecules, which act as signals (4). In Gram-negative bacteria, these are AHLs, which are often synthesised by LuxI-type synthases and recognised by LuxR-type receptors (5). In the case of some Enterobacteria, such as *K. pneumoniae* and *Escherichia coli*, a putative LuxI synthase is absent, yet an orphan LuxR receptor (SdiA) is present and it has been shown that this receptor is capable of detecting AHLs produced by other bacteria (6).

The regulatory role of SdiA in virulence has been widely investigated in *E. coli*. It has been demonstrated that SdiA has a promoting effect on survival in the gastrointestinal tract through acid tolerance upregulating *gad* expression (7), and the promotion of resistance to quinolones through expression of AcrAB efflux pump (8). Conversely, SdiA has also been demonstrated to exert a repressive effect on other phenotypes, including motility and adhesion, through repression of *fliC* (flagella) and *fimA* (fimbriae) expression (6). Furthermore, SdiA has been proposed as a repressor of biofilm formation, as SdiA-lacking strains show lower biofilm forming ability through *uvrY* repression (9). Nevertheless, there are also studies that argue that SdiA does not affect biofilm formation in this species (7). However, the majority of laboratory strains of *E. coli* are low biofilm formers, which may result in an underestimation of the impact of QS on this phenotype (10). Furthermore, a study conducted with a SdiA-lacking *Cronobacter sakazakii* strain showed increased expression of capsule and lipopolysaccharide (LPS) synthesis genes (11), and other research has highlighted the importance of SdiA in conferring resistance to serum and complement killing in *Salmonella* (12).

SdiA has been shown to perform regulatory functions in an AHL-dependent manner. In *E. coli*, some studies observed that the exogenous addition of various AHLs reduces biofilm formation through the binding of SdiA (13). Moreover, an increased sensitivity to phage infection was observed in *E. coli* following treatment with AHLs in the presence of SdiA (14), indicating that SdiA plays a role in bacteriophage sensitivity through an AHL-dependent mechanism.

Nevertheless, to date, no native SdiA ligand has been identified in *E. coli*, as some studies have indicated that SdiA may have a promiscuous role, as this receptor has been shown to bind to a diverse range of ligands, including AHLs and synthetic molecules (15,16).

The role of QS signalling in *K. pneumoniae* has only been the subject of a limited number of studies, and there is considerable inconsistency in the literature regarding AHL-regulated QS. For instance, some authors have proposed that *K. pneumoniae* is devoid of the *luxI* homologues for synthesis of AHLs (17,18). However, other researchers have reported the production of AHLs in *K. pneumoniae* strains (19–21). With regard to SdiA, in the study conducted by Pacheco et al. (2021) proposed that SdiA functions as a repressor of biofilm formation and fimbriae expression in *K. pneumoniae*. To conduct the study, the authors employed a *sdiA* transposon-based insertion mutant of the *K. pneumoniae* ATCC 10031 strain and examined the impact of *N*-octanoyl-L-homoserine lactone (C8-HSL) as an exogenous AHL on a microtiter-based biofilm formation cultivation model. In this study, the authors observed that the biofilm-repressing effect of the AHL was SdiA-dependent.

The aim of this study is to elucidate the function of SdiA and AHL supplementation in virulence-related traits and biofilm formation of *K. pneumoniae*. Virulence phenotypes, including biofilm formation, capsular synthesis, serum resistance, and phage sensitivity, was evaluated, plus virulence assessment in vivo in *Galleria mellonella*. The *K. pneumoniae* strain selected for this study was KLEB-33, a convergent, multiresistant, hypermucoviscous and hyperbiofilm-forming strain with multiple hypervirulence genes (22). The genetic and phenotypic characteristics of KLEB-33 render it an optimal model for the study of the emerging convergent *K. pneumoniae* strains. For comparative purposes, the aforementioned phenotypes were also studied in a non-virulent, low biofilm forming, and non-MDR ATCC 13883^T^ strain. The results of this study demonstrate that C6-HSL and SdiA exert a dual influence on virulence phenotypes, operating both independently and hierarchically. The experiments conducted have facilitated a more profound comprehension of the QS mechanisms in *K. pneumoniae*.

## 4 Materials and Methods

### 4.1 Bacterial strains and culture conditions

This study employed the *K. pneumoniae* ATCC 13883^T^ and KLEB-33 strains. KLEB-33 is a multiresistant hyper-biofilm-forming strain harbouring hypervirulence genes (22). The strains were routinely grown at 37 °C/200 rpm on 5 mL Lysogeny broth (LB) or LB agar (1.5 % w/v). Antibiotics were added when required, and synthetic AHL signals dissolved in acetonitrile at a concentration of 10 g/L, and comprising acyl chains with a carbon length of 4 to 18, were added at a final concentration of 5, 2 or 0.2 µM, as required.

### 4.2 Construction of *sdiA* mutants

The *sdiA* gene was deleted from the ATCC 13883^T^ and KLEB-33 strains using the CRISPR/Cas9-based system with the pCas9KP-Apr and pSGKP-Km plasmids, as previously described (23). The sequences of the single-guide RNA (sgRNA) spacer, the single-stranded DNA (ssDNA) sequences employed for allelic knockout, and the primers used for mutation confirmation are shown in **Table S1**. Completely removal of the gene was confirmed by Sanger sequencing.

Planktonic growth was assessed in 15 mL LB cultures, plus the effect of AHL addition. Briefly, cultures inoculated at an initial absorbance at 600 nm of 0.01 (Abs_600 nm_) were incubated at 200 rpm, 37 °C/24 h, with growth measured at 1, 2, 4, 6, 8, 10 and 24 h, in triplicate. The lineal relationship between Colony Forming Units (CFUs) and Abs_600 nm_ was confirmed using the Miles and Misra method (24), with the experiments being repeated twice.

### 4.3 Biofilm cultivation

1. *K. pneumoniae* KLEB-33 biofilms were cultivated in LB using the active attachment (AA) method as previously described (25). Briefly, biofilms were grown for 24 h/37 °C in 12-well plates (VWR, 734-2778) using a custom-made aluminium lid with glass coverslips (18x18 mm) attached as a substrate. Bacteria were inoculated at final Abs_600 nm_ of 0.05. The culture media and treatment were replaced at 12 h to facilitate the growth of adherent cells. The Rolling Biofilm Bioreactor (RBB) cultivation method (26) was also used to promote biofilm growth and maturation of KLEB-33 and ATCC 13883^T^ strains of *K. pneumoniae*. In this system bacteria were inoculated at a final Abs_600 nm_ of 0.01 and incubated at 37 °C/72 h, with media and treatment changes every 24 h. The biofilm biomass was quantified by staining with crystal violet (0.04 %) and measuring the absorbance at Abs_590 nm_ after washing the coverslips with 33 % acetic acid (27).

RBB biofilms were stained with Syto9 (ThermoFisher S34854) and subsequently imaged by confocal laser scanning microscopy (CLSM) (Leica Stellaris 8) to quantify biofilm height and coverage at 24, 48 and 72 h. Furthermore, 24 h biofilms were also examined for bacterial filamentation. To examine the composition and structure of biofilms, 48 h samples were also stained with YOYO™-1 iodide (ThermoFisher Y3601), Concanavalin A conjugated with Alexa Fluor® 594 (ThermoFisher C11253) and lipophilic FM™ 4-64 (ThermoFisher, F34653) fluorescent dyes to stain biofilm extracellular DNA (eDNA), extracellular polysaccharides, and cell membranes, respectively. A total of six fields were collected per sample. Images were subsequently analysed using ImageJ (v1.54) and Leica Application Suite X Office (v1.4.6.28433).

### 4.4 Quorum Quenching activity

Quorum quenching (QQ) activity was evaluated in accordance with the methodology previously described (28). Briefly, 500 µL of the filtrated (0.22 µm) supernatant and pellet (resuspended in PBS pH 6.5) portions of 15 mL 24 h LB culture samples were exposed to C6-HSL (10 µM) for 6, 12, 24 and 48 h. pH of samples was adjusted when necessary to 6.5 to prevent the spontaneous opening of the lactone ring. PBS pH 6.5 with C6-HSL (10 µM) was used as negative control.

After incubation, 100 µL of each sample was added to wells prepared in soft LB agar plates (0.8 %) with the biosensor *Chromobacterium subtsugae* CV026, and incubated at 30 °C/24 h. Pellet samples were filtrated to avoid contamination of biosensor. Absence of production of violacein by the biosensor was indicative of positive QQ activity. Biosensor was routinely grown in LB broth supplemented with kanamycin (25 µg/mL).

### 4.5 Percoll density gradient centrifugation and Capsule staining

Percoll density gradient centrifugation was employed to quantify strain capsule expression, in accordance with the methodology described (29). Bacteria were adjusted to Abs_590 nm_ = 1, collected from overnight cultures by centrifugation and resuspended in 2 mL PBS. The bacterial suspension was added to the top of a Percoll density gradient comprising 80, 60, 40 and 20 % solutions in PBS to separate the bacterial fractions after centrifugation (2600 g) at 4 °C/20 min (9x acceleration; 1x deceleration). The distance between the bottom of the tube and the cell layer was measured. Capsule staining was also conducted using the Maneval method (30).

### 4.6 Human serum sensitivity

The susceptibility to human serum was assessed as previously described (29,31). Briefly, the bacterial inoculum was adjusted to an Abs_600 nm_ of 1 in PBS, and 100 µL of bacterial suspension was added to 200 µL of pre-warmed (37 °C) human serum (Merk, S7023). Mixture was incubated at 37 °C/2 h. Colony-forming units per millilitre (CFU/mL) were determined on LB agar plates.

### 4.7 Phage susceptibility

Phage susceptibility was evaluated using the specific lytic phage *Webervirus kpv33d1* (Sonia Rey et al., unpublished) as previously described (32). Briefly, the bacterial inoculum was grown in LB to an Abs_600 nm_ of 0.5, diluted 1/5, and 10 µL were added to each well containing 90 µL of pre-diluted phage, resulting in a final Abs_600 nm_ of 0.01 (approximately 10^6^ CFU/mL). Phage at 10^10^ Plate Forming Units (PFU)/mL was serially diluted 1/10 to up to 9 times in 90 µL LB in 96-well U-bottom plates, from a Multiplicity Of Infection (MOI) of 10^3^ to 10^-5^. A gas-permeable membrane (Breathe-Easy®, Z380059) was applied, and the plate was incubated at 37 °C/24 h, with Abs_600 nm_ readings of the cultures every 30 min.

### 4.8 Galleria mellonella infection model

The virulence was evaluated using the *G. mellonella* survival assay, as previously described (33,34). Briefly, 15 larvae (300 - 400 mg body weight) were injected with 10 μL of a suspension containing 10^3^, 10^5^, or 10^7^ CFUs in PBS. Larvae injected with an equal volume of sterile PBS or PBS plus C6-HSL (5 µM) were used as controls. Larvae were incubated at 37 °C in the dark and mortality was monitored every 24 h for up to 3 days.

### 4.9 Statistical analysis

Statistical analyses were conducted using GraphPad Prism 8.3.0. First, a Shapiro-Wilk test was used to ascertain whether the data exhibited a normal distribution. If normal, an analysis of variance (ANOVA) or a Student’s t-test was conducted. Alternatively, a Kruskal-Wallis or a Mann-Whitney test was performed, depending on whether there were more than two groups or only two groups, respectively. The significance values indicated by asterisks in the graphs presented in this paper are as follows: * = p<0.05; ** = p<0.005; *** = p<0.0005; and **** = p<0.00005.

## 5 Results and Discussion

### 5.1 C6-HSL exhibited the greatest effect in promoting biofilm formation in AA cultivation system

Prior research has shown that SdiA exhibits a high degree of promiscuity, as evidenced by its ability to recognize different ligands (15). Another study has indicated that SdiA contains a range of amino acids within its active site that are capable of interacting with up to ten different AHLs (35). It is therefore necessary to perform a screening of different AHLs in a robust and repeatable biofilm cultivation method to elucidate the function of SdiA in biofilm formation following AHL addition in *K. pneumoniae*. To this end, the effect of three short-chain homoserine lactones (C4-HSL, C6-HSL and C8-HSL) and five long-chain homoserine lactones (C10-HSL, C12-HSL, C14-HSL, C16-HSL and C18-HSL) on the AA biofilm cultivation system in the KLEB-33 strain was investigated. The AA system has been already demonstrated to be a reliable and repeatable system for biofilm studies in this strain (25). The results showed that the AHLs C4-HSL and C6-HSL had the most pronounced impact by increasing biofilm formation at 5 µM (**Figure 1A**), with no effect at 2 µM. With C12-HSL and C14-HSL also exhibiting a significant effect, albeit with a considerably lower magnitude than that observed for the short-chain AHLs (**Figure 1B**).

**Figure 1.**
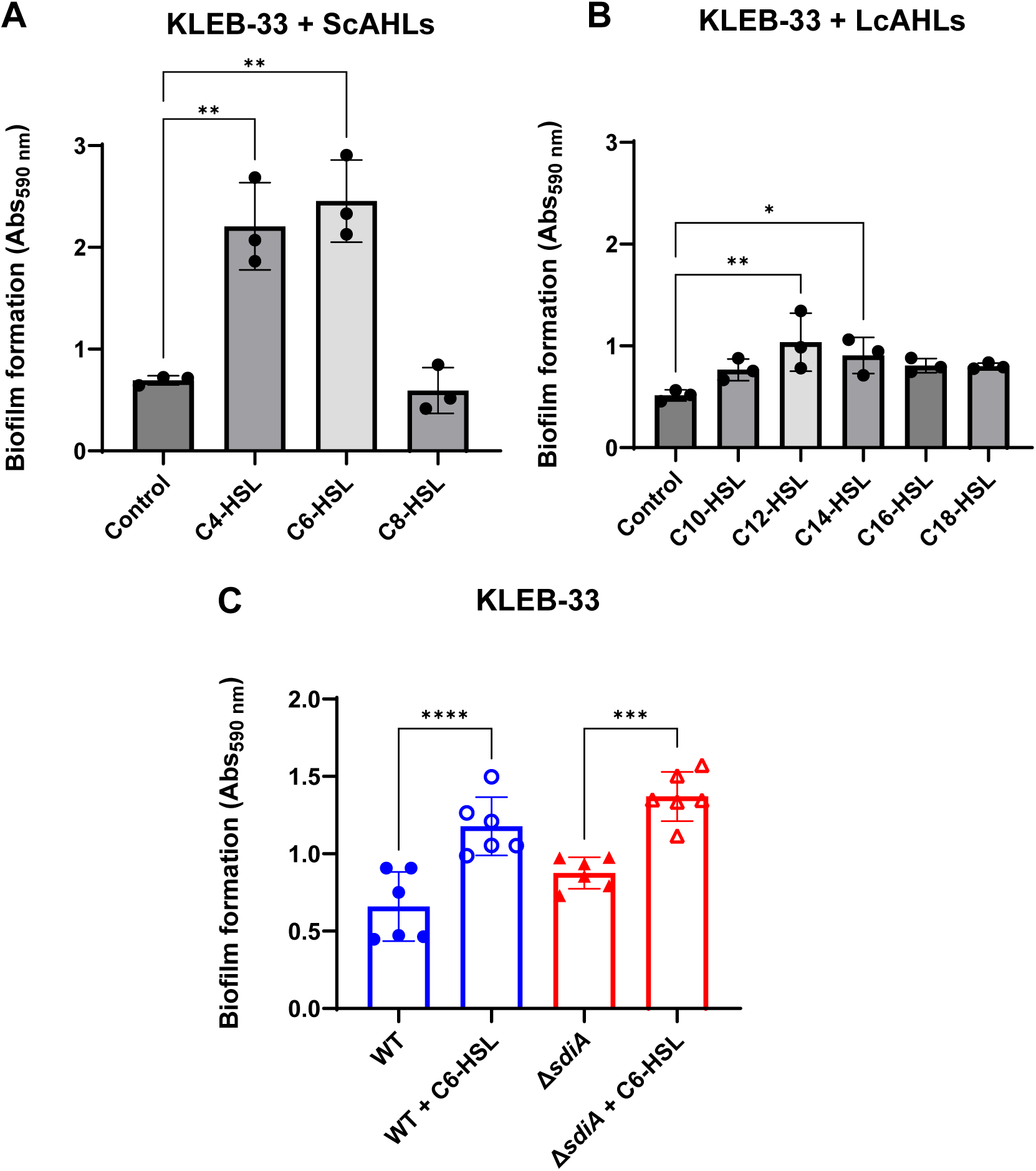
Effect of AHL addition (5 µM) on biofilm formation in *K. pneumoniae* KLEB-33. The quantification of biofilm formation was conducted using Crystal Violet (CV) staining, which was subsequently dissolved with 33 % acetic acid and the absorbance measured at 590 nm (Abs_590 nm_). No effect was observed at 2 µM in previous experiments (data not shown). The impact of each AHL individually at 5 µM was examined for short-chain AHLs (ScAHLs) (**A**) and long-chain AHLs (LcAHLs) (**B**). The repeatability of the C6-HSL effect was validated in subsequent experiments conducted on wild-type and SdiA-deficient KLEB-33 strains (**C**). All the experiments were conducted in duplicate.

In contrast with observations made by Pacheco et al. (2021), our findings did not indicate a promoting effect on biofilm formation of C8-HSL. This suggests that this AHL may not exert a physiological effect on the strain studied. Given the pronounced biofilm-promoting impact of C6-HSL observed in our experiments, we sought to ascertain whether this effect could be attributed to its direct interaction with SdiA. To this end, a Δ*sdiA* strain was constructed in KLEB-33 and was cultivated under C6-HSL supplementation. A significant increase in biofilm formation was also observed in the SdiA-lacking strain stimulated by C6-HSL addition (**Figure 1C**), indicating that the biofilm-promoting effect of C6-HSL was independent of SdiA. Moreover, despite SdiA being described in the literature as a biofilm repressor (6,17), our experiments revealed only a slight increase in biofilm formation on the Δ*sdiA* in comparison to its wild-type in the AA biofilm cultivation system. Growth was monitored with/without AHL supplementation in shaken cultures to ascertain that SdiA deficiency and/or C6-HSL addition does not affect growth (**Figure S1**).

It is noteworthy that the concentration of C6-HSL at which a significant impact on biofilm formation was observed is higher than the physiological AHL concentrations typically encountered in QS signalling species (36). Nevertheless, the same minimal concentration has been necessary to elicit a biological effect in *K. pneumoniae* before (37). This could be due to the QQ activity present in this species, as AHL-degrading enzymes have previously been described (38), and the presence of QQ activity against C6-HSL was corroborated in the culture media of both strains (**Table S2**). The influence of C6-HSL provision on ATCC 13883^T^ strain cultures was also assessed in the AA cultivation system. However, the inherent low biofilm-forming capacity of the strain (25) hindered the detection of significant differences in this biofilm model system.

### 5.2 The formation and maturation of *K. pneumoniae* biofilms are influenced by SdiA and C6-HSL in an opposing and hierarchically organised manner

In order to evaluate the impact of *sdiA* mutation and C6-HSL supplementation on biofilm structure and maturation, we also employed the rolling biofilm bioreactor (RBB) system (26). This system permits the cultivation of biofilms over extended periods and the generation of highly matured biofilms with high reproducibility, even in strains with low biofilm-forming capabilities, such as ATCC 13883^T^. The results obtained with the RBB system demonstrated that the KLEB-33 Δ*sdiA* strain exhibited a higher biofilm maturation in comparison to the wild-typestrain. This was evidenced by the earlier formation of mushroom-like structures in the Δ*sdiA* strain after 48 hours of incubation (**Figure 2**). Our observations in the RBB cultivation system are in accordance with a role of SdiA as a biofilm-repressor, as previously described in the literature (6,17). Moreover, no notable differences were observed between the wild-type and Δ*sdiA* strains at 24 hours of incubation, as happened in the AA cultivation system, as the 24-hour biofilms had not yet reached a stage of development sufficient to manifest such differences.

**Figure 2.**
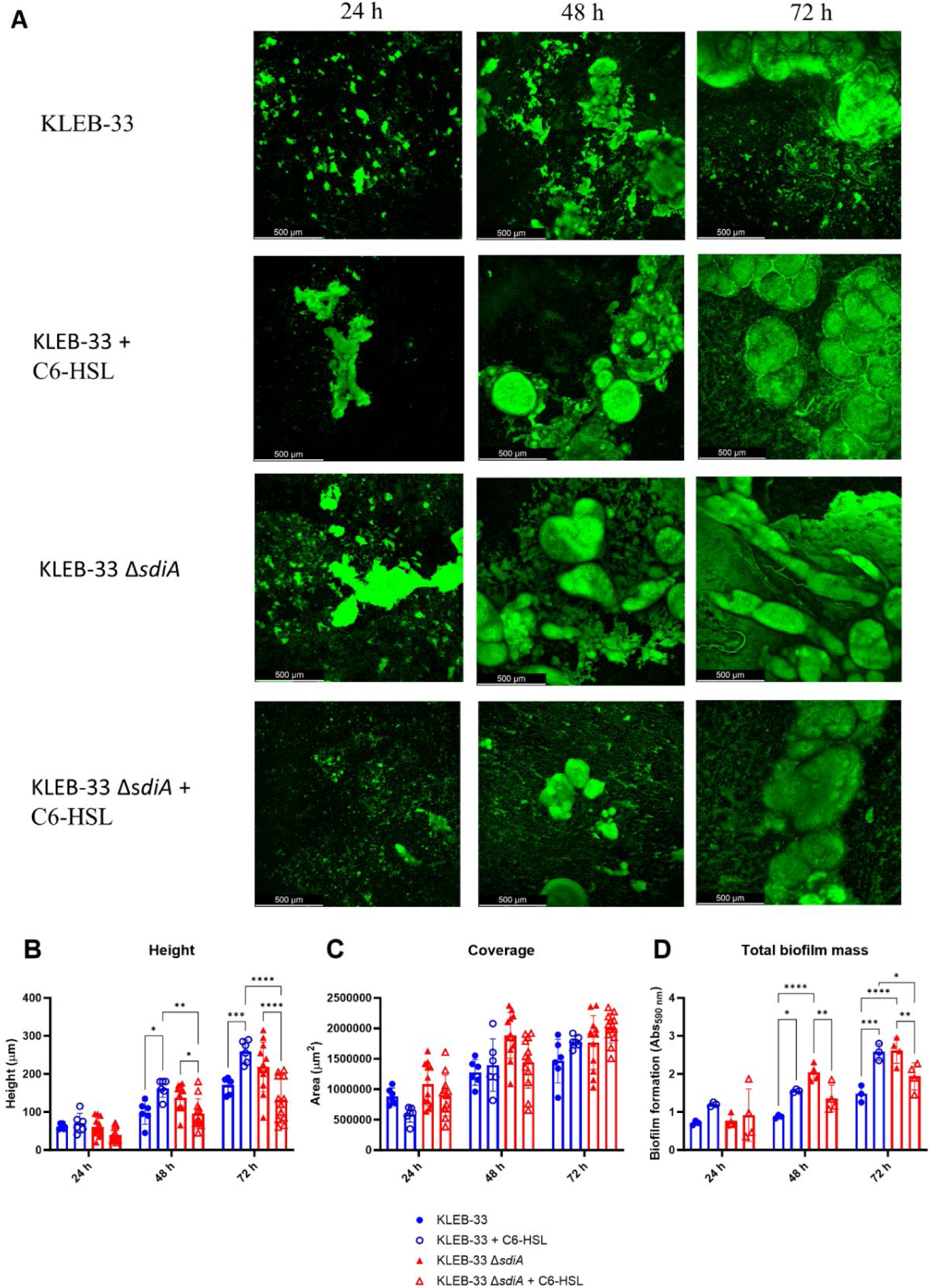
Impact of *sdiA* mutation and AHL addition on biofilm formation in *K. pneumoniae* KLEB-33. (**A**) Representative CLSM images of biofilms obtained after 24, 48 and 72 hours of incubation in the RBB cultivation system and staining with Syto9 fluorescent dye. (**B**) and (**C**) Quantification of height and biofilm coverage from confocal images using ImageJ (v1.54) image analysis software. (**D**) Crystal Violet quantification of wild-type and Δ*sdiA K. pneumoniae* KLEB-33 strains biofilm biomass after treatment with C6-HSL (5 µM).

Indeed, comparable levels of biofilm formation were recorded in the AA and RBB cultivation systems for both strains following a 24-hour incubation period (**Figure 1** and **Figure 2**).

Nevertheless, no significant increase in biofilm formation was observed in the ATCC 13883^T^ Δ*sdiA* strain. This finding may again be attributed to the inherent lower biofilm formation ability of the strain. However, the data on biofilm height showed a notable elevation in the mutant strain, indicative of higher maturation of the biofilm communities (**Figure S2**).

The addition of C6-HSL (5 μM) also promoted the maturation of the biofilm in the KLEB-33 and ATCC 13883^T^ wild-type strains after 48 hours of incubation (**Figure 2** and **Figure S2**). However, no such promotion effect was observed in the Δ*sdiA* strain when AHL was added for both strains studied, and these observations were corroborated by the quantification of the total biofilm biomass and thickness (**Figure S2** and **Figure 2**). These results show that in the wild-type strain, SdiA functions as a repressor of genes involved in biofilm maturation. However, this effect is negated in the presence of AHL, resulting in a biofilm-promoting effect only when SdiA is present. Interestingly, in the mutant strain, while the absence of SdiA promotes biofilm maturation, this is diminished in the presence of C6-HSL. This indicates the potential involvement of genes that regulate biofilm growth and are repressed by C6-HSL signalling when SdiA is absent. The observed phenotype is only evident in the absence of SdiA, suggesting a hierarchical regulatory relationship between SdiA and C6-HSL.

Additionally, a significant increase in cell filamentation was observed in biofilms of the KLEB-33 Δ*sdiA*, a finding that aligns with previous observations (17) (**Figure 3** and **Figure S3**). The formation of filamented bacteria has been documented in the literature as a mechanism that contributes to the persistence and colonisation of surfaces (39). Therefore, the elevated filamentation rates observed in the Δ*sdiA* strain are consistent with its enhanced capacity for biofilm formation (**Figure 2**). However, no differences were observed following C6-HSL supplementation, indicating that cell filamentation is dependent on SdiA, yet not triggered by AHL signalling. This provides further evidence for the existence of a separate regulatory mechanism.

**Figure 3.**
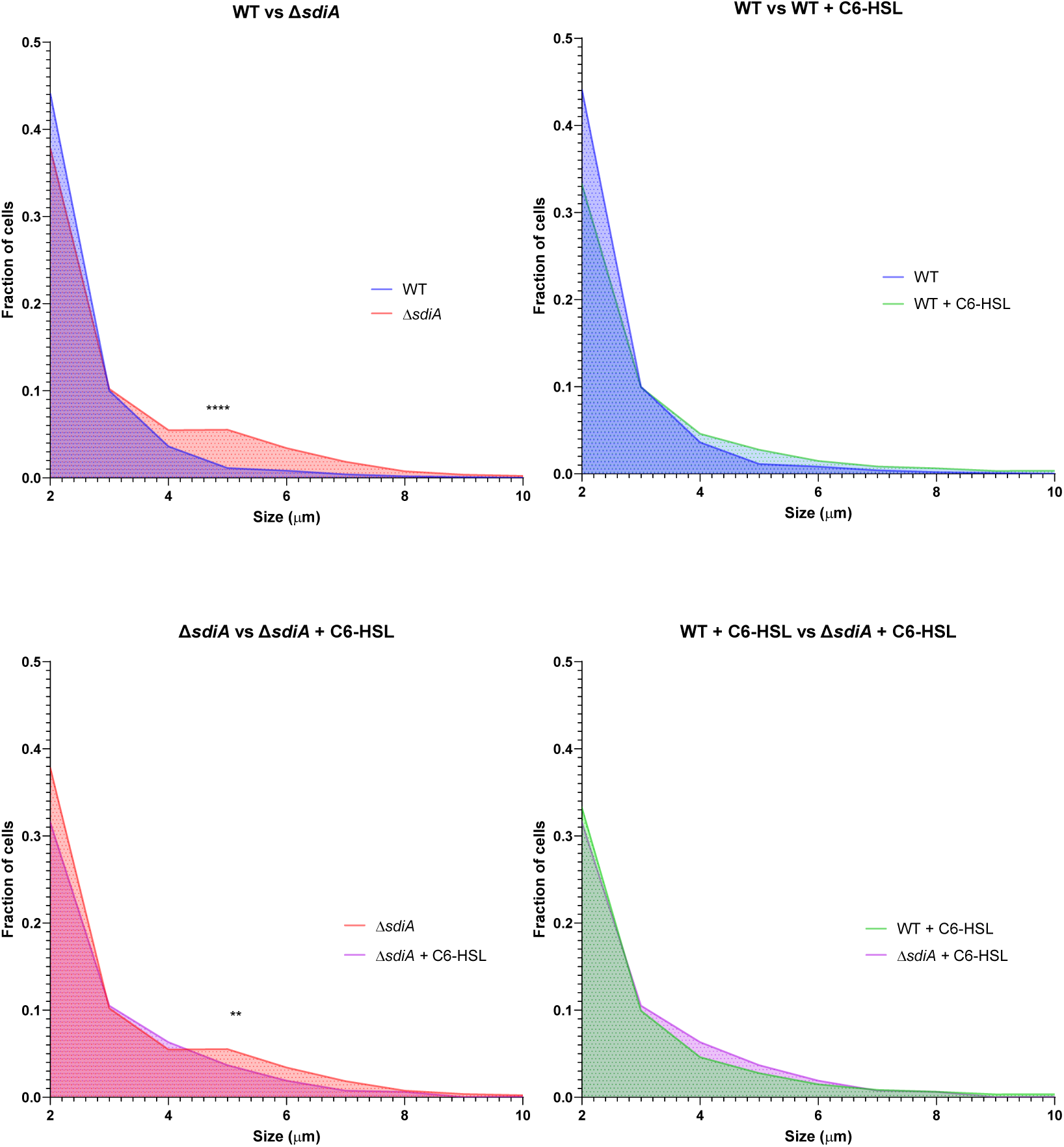
Distribution of cell size fractions of *K. pneumoniae* KLEB-33 wild-type (WT) and Δ*sdiA* strains biofilms cultivated in the RBB system for 24 hours with and without C6-HSL supplementation (5 µM). Cell sizes were measured using the ImageJ software (version 1.54).

In light of the observed alterations in biofilm maturation and structure following *sdiA* mutation and AHL supplementation, we sought to investigate whether SdiA deficiency or AHL signalling could influence the biofilm matrix composition in *K. pneumoniae*, as previously described for other pathogens (40,41). To this end, 48-hour RBB biofilms of KLEB-33 were fluorescently stained with YOYO-1 (eDNA), FM 4-64 (cell membranes) and Concanavalin A conjugated with AlexaFluor594 (ConA, extracellular polysaccharides). The results demonstrated a significant increase in fluorescence intensity for both the YOYO-1 and FM 4-64 signals (**Figure 4**) in the same conditions where enhanced biofilm formation was recorded (**Figure 2**: WT + C6-HSL and Δ*sdiA*). However, the ConA staining did not yield a strong fluorescence signal, which could suggest that eDNA may be the principal component of the biofilm matrix (**Figure S4**). The function of eDNA as a crucial component of the biofilm matrix, facilitating the development of biofilms through the formation of adhesive "webs" (**Figure S4**) that enhance cell cohesion and adhesion, has been previously postulated in a variety of bacteria (26,42,43). However, no clear relationship could be identified between SdiA or AHL addition and eDNA. This suggests that the regulation of this matrix component is not dependent on this regulatory system.

**Figure 4.**
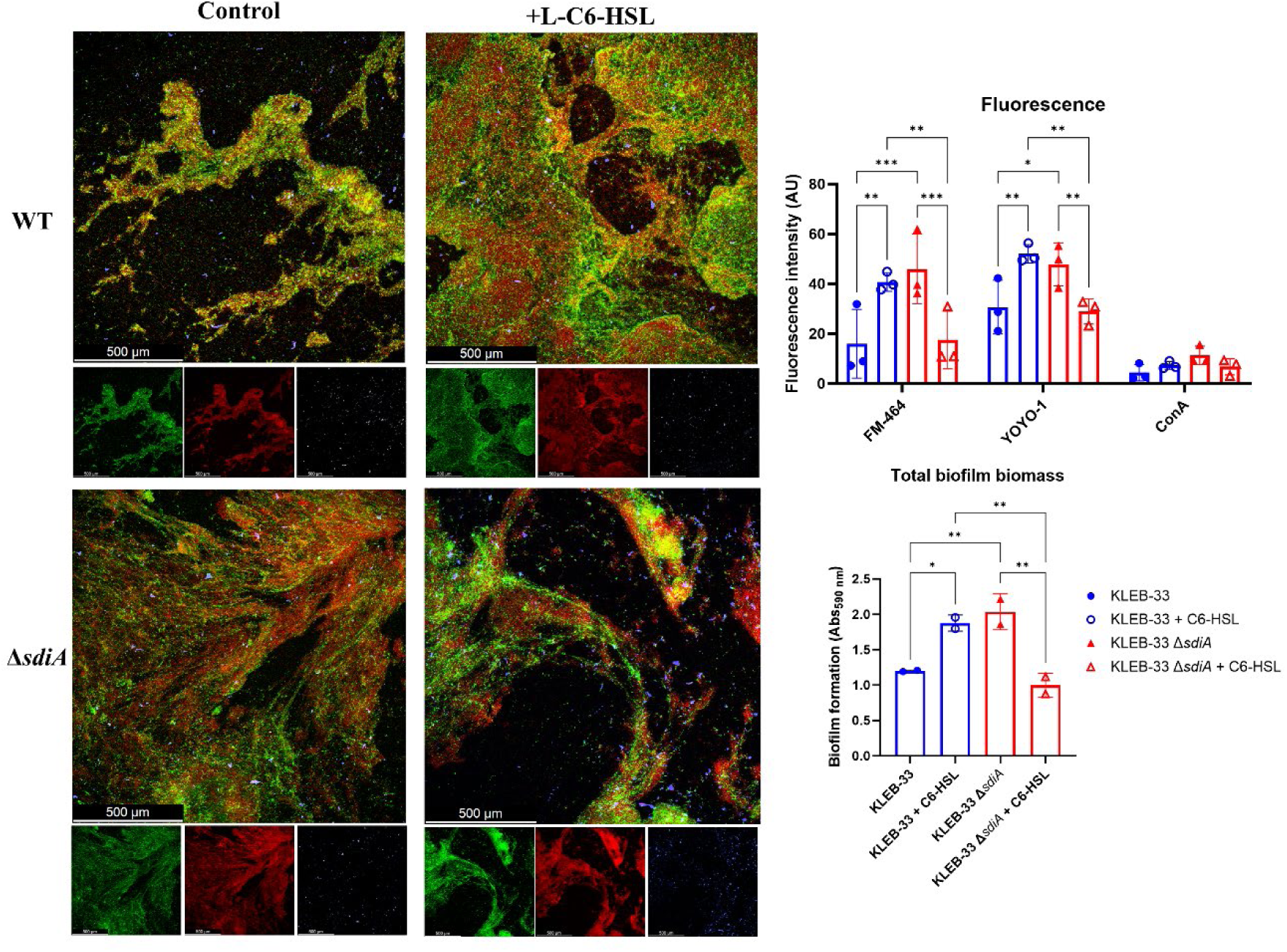
Representative CLSM (Leica Stellaris 8) images of RBB biofilms of WT and Δ*sdiA K. pneumoniae* KLEB-33 48 h treated with C6-HSL (5 µM). Biofilms were stained with YOYO-1 (eDNA, green), FM-464 (cell membranes, red), and Concanavalin A conjugated with AlexaFluor 594 (exopolysaccharide, blue). A total of tree fields were recorded in CLSM for each biofilm sample. Repeatability of the experiments was corroborated with total biomass quantification with CV staining (Abs_590nm_). Quantification of fluorescence intensity (AU: arbitrary units) was performed with ImageJ (v1.54).

### 5.3 QS-induced changes in biofilm formation have no correlation with capsule production in *K. pneumoniae*

To further examine the mechanism underlying the alterations in biofilm formation observed, we postulate that the supplementation of C6-HSL or the mutation of *sdiA* (which is associated with high biofilm formation) may be related to a reduction in capsule production. This is based on the findings of other researchers who have demonstrated that high-capsule-producing bacteria are more likely to be low biofilm formers, as the capsule polysaccharides have been shown to interfere with adhesion (44). Consequently, a semi-quantitative analysis of the capsule production was conducted using Percoll density gradient centrifugation. This method enables the macroscopic differentiation of high and low capsule-producing bacteria based on their flotation characteristics (29). Additionally, the capsules of the strains under study were subjected to staining using the Maneval’s method (30). The results revealed no appreciable differences in capsule production between the wild-type and the Δ*sdiA* for both strains studied (**Figure S5** and **Figure 5**). Regarding the addition of C6-HSL, a significant increase in capsule production was recorded in the wild-type strain of ATCC 13883^T^, whereas no effect was observed in KLEB-33 floatability (**Figure 5**), which is likely attributable to its relatively lower capsule production compared to the ATCC 13883^T^ strain (**Figure S5** and **Figure 5**). These findings indicate that the promotion of capsule biosynthesis by C6-HSL supplementation is dependent on SdiA, as no effect was observed in a strain of ATCC 13883^T^ lacking this receptor. Nevertheless, our results also suggest that there is no direct correlation between capsule production and biofilm formation in the strains under investigation. This is evidenced by the fact that the conditions that promoted biofilm formation, namely the addition of C6-HSL to wild-type strains or *sdiA* mutation, did not result in a reduction in capsule production. Moreover, and although other researchers have reported elevated polysaccharide expression in a SdiA-deficient *C. sakazakii* strain (11), no changes in exopolysaccharide abundance were observed in CLSM biofilms images (**Figure 3**).

**Figure 5.**
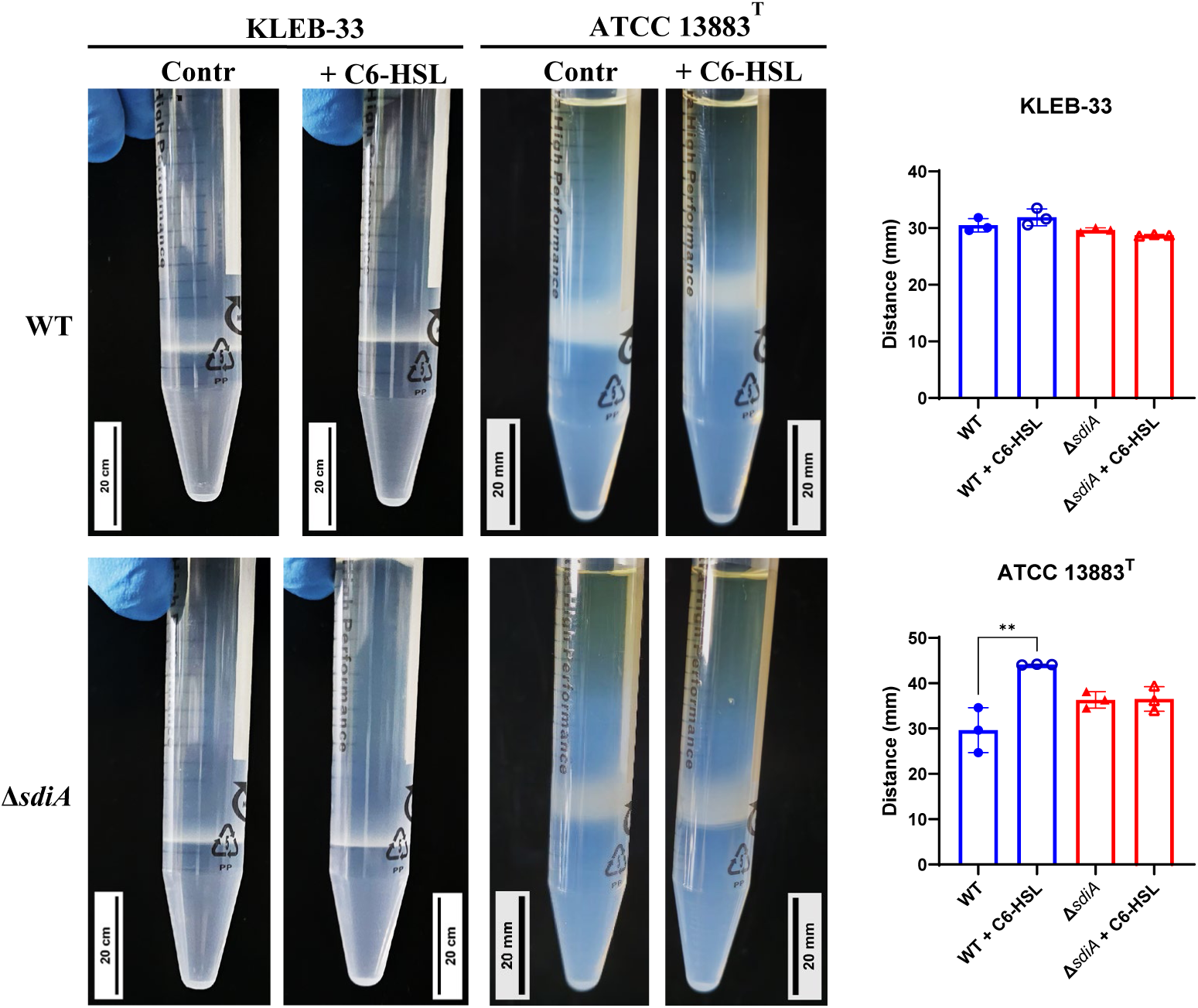
Percoll density gradient analysis conducted on *K. pneumoniae* KLEB-33 and ATCC 13883^T^ wild-type (WT) and *ΔsdiA* strains with and without the addition of AHL (5 µM). Representative images are presented in (**A**), while histograms of the obtained measurements for bacterial cell layer height are shown in (**B**).

### 5.4 Serum complement killing in *K. pneumoniae* is inversely affected by SdiA and AHL supplementation

Although the Percoll method and Maneval’s staining did not reveal macroscopic differences in capsule production between strains, alterations in capsule composition may underlie the observed variations in biofilm formation. Consequently, we sought to identify a phenotype that could depend on capsule composition and be sufficiently sensitive to detect differences in capsule biosynthesis. To this end, we conducted human serum survival assays as this method is employed by some authors as an indirect means of assessing capsule production (31). This is because capsule biosynthesis has been associated with serum and complement-killing resistance (2). The results of our experiments showed that the *sdiA* knockout strains exhibited significantly elevated serum sensitivity in both strains under study (**Figure 6**), thereby indicating that SdiA plays a role in this process.

**Figure 6.**
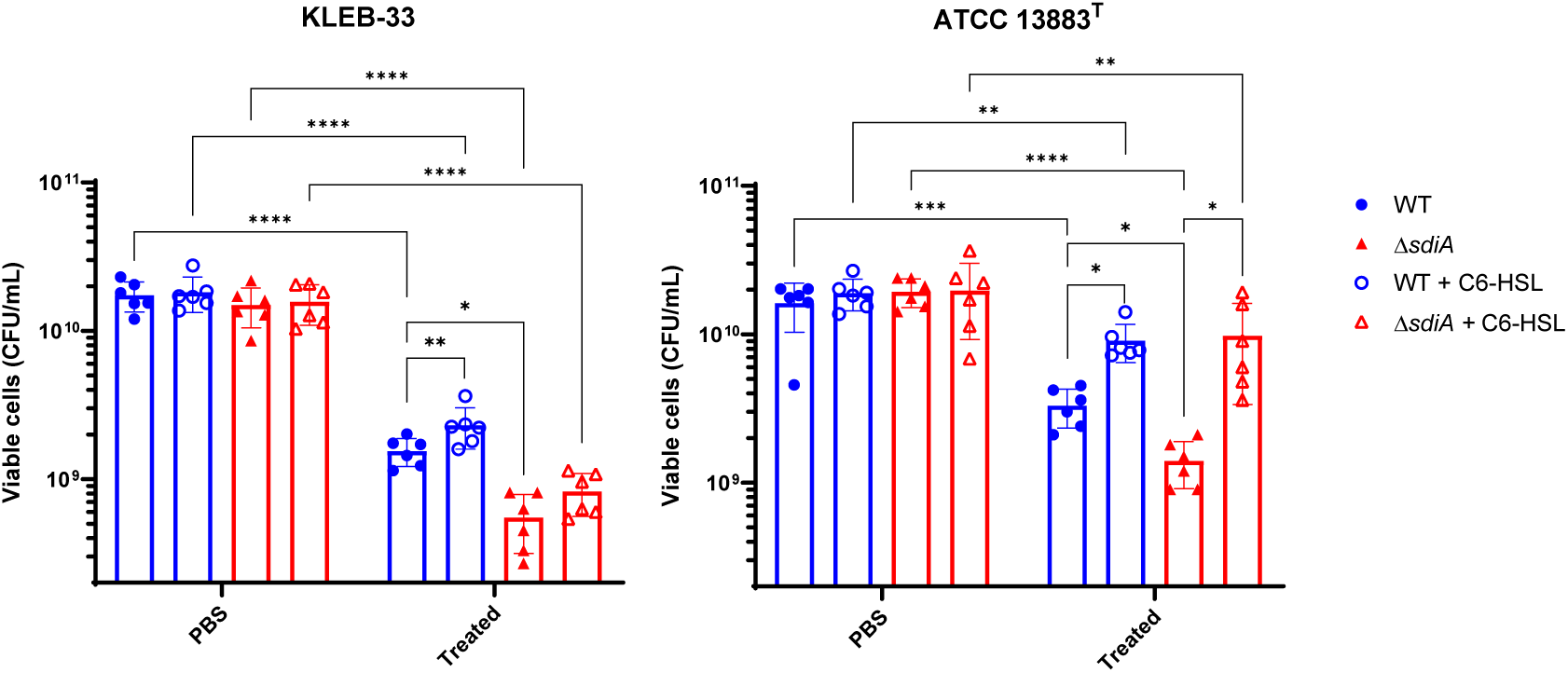
Serum resistance analysis of *K. pneumoniae* KLEB-33 (**A**) and ATCC 13883^T^ (**B**) wild-type (WT) and Δ*sdiA* strains following exposure to human serum for 2 hours, plus the effect of C6-HSL supplementation (5 µM). The control samples represent non-serum-treated bacteria in PBS.

The reduced serum resistance exhibited by the Δ*sdiA* strains may be linked to modifications in capsule production, which could also account for the increased biofilm formation. However, an alternative hypothesis is that an SdiA deficiency in *K. pneumoniae* may lead to alterations in other components of the cell surface that affect serum resistance. For instance, a number of studies indicates that LPS upregulation contributes to higher serum and phage sensitivity (2,45,46). Moreover, the diminished serum resistance observed in the Δ*sdiA* strains is consistent with the findings of previous studies conducted in *Salmonella* spp, as SdiA enhances serum resistance through the expression of *rck*, an open reading frame encoding an outer membrane protein associated with complement-mediated killing resistance (12). Additionally, type-1 fimbriae were found to be overexpressed in a SdiA-deficient strain of *K. pneumoniae* (17), and its overexpression has been linked to detrimental effects on complement survival in *E. coli* (47).

Conversely, the addition of C6-HSL was observed to increase serum resistance independently of the presence of SdiA (**Figure 6**). Therefore, the addition of C6-HSL does not appear to be associated with reduced capsule production, as evidenced by the observation of enhanced serum resistance, a heightened band in Percoll experiments in the ATCC 13883^T^ strain, and elevated biofilm formation.

### 5.5 SdiA deficiency increases phage sensitivity in *K. pneumoniae*

The quantification of capsule production and complement-killing resistance experiments suggested that SdiA may affect cell surface components other than capsule, independently of C6-HSL. Furthermore, a study conducted in *E. coli* demonstrated that SdiA plays a role in bacteriophage sensitivity through an AHL-dependent mechanism (14). Accordingly, an experiment was conducted to investigate whether modifications to the cell surface resulting from QS signalling could influence the susceptibility to phage infection. The infectivity of *Webervirus kpv33d1*, a high lytic wastewater-derived phage isolated against KLEB-33 strain, was tested on the genetically modified strains lacking the *sdiA* gene with or without AHL supplementation. The results showed a dose-response relationship with regard to the phage dose and differences in phage multiplicity of infection (MOI) attributable to host specificity (**Figure 7**). The deletion of the *sdiA* gene resulted in a growth inhibition only in the mutant strains in the presence of a phage MOI of 10 and 0.001 in the KLEB-33 and ATCC 13883^T^ strains, respectively. The addition of C6-HSL at concentrations of 5, 2 and 0.2 µM did not result in any observable effect on phage susceptibility (data not shown). These findings are consistent with those previously observed in *E. coli* (14) and also are in align with the results of serum resistance experiments, as evidenced by the heightened sensitivity of the Δ*sdiA* strains compared to the parental strains. This may be attributed to an elevated expression of LPS, which has been previously documented in a Δ*sdiA* strain of *C. sakazakii* (11).The overproduction of LPS has been demonstrated to contribute to higher sensitivity to phages, due to its recognition as a bacterial epitope (46). The observation of higher serum and phage sensitivity of Δ*sdiA* may alternatively be related to the higher filamentation rates observed, with higher cell surface being exposed to complement killing and phage attachment. However, the fact that C6-HSL addition has no effect on phage infection provides further support for a regulatory role of SdiA independent of C6-HSL.

**Figure 7.**
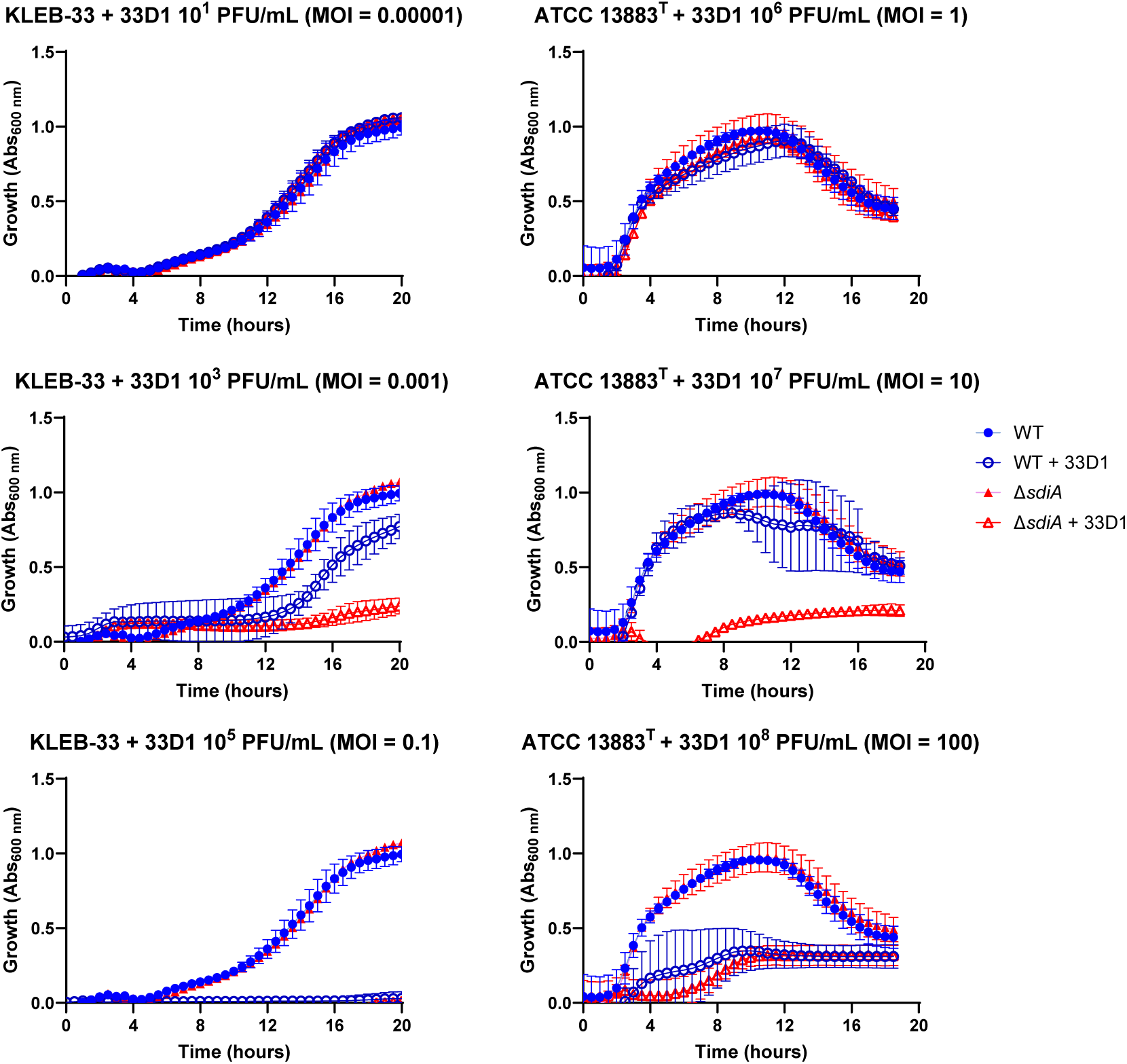
Susceptibility assays of *K. pneumoniae* KLEB-33 and ATCC 13883^T^ wild-type (WT) and Δ*sdiA* strains (10^6^ CFU/mL) to *Webervirus kpv33d1* phage. A dose-response assay was conducted using 96-well microtiter plates and varying Multiplicity Of Infection (MOI) values. Experiments were performed for each strain separately. No effect of C6-HSL addition (5 µM) was observed (data not shown).

### 5.6 C6-HSL signalling increases *Galleria mellonella* survival after infection with SdiA-deficient *K. pneumoniae*

Our earlier findings suggested that mutation of SdiA could be responsible for alterations in the surface characteristics of *K. pneumoniae* cells and enhanced biofilm maturation. A previous study has demonstrated overexpression of type-1 fimbriae in a *sdiA* mutant of *K. pneumoniae*, which could be a contributing factor to higher biofilm formation (17). This provides a potential explanation for our observations, but it could also be a contributing factor to higher virulence and mortality rates *in vivo* (2). To test this, we examined whether strains lacking SdiA could exhibit higher mortality rates in the *G. mellonella* infection model. As a result, despite the mortality rates recorded daily being consistently higher in the mutant, no statistically significant differences were observed between the wild-type and *ΔsdiA* strains for both KLEB-33 and ATCC 13883^T^ (**Figure 8A**). Regarding the addition of C6-HSL, a markedly reduced virulence was observed in the KLEB-33 *ΔsdiA* strain at a dose of 10^5^ CFU in the presence of AHL in comparison with the absence of AHL supplementation (**Figure 8B**). However, C6-HSL had no effect on virulence in either of the wild-type strains. The effect of decreased virulence observed following AHL treatment of the *sdiA* mutant strain was not replicated in ATCC 13883^T^. This may be attributed to the higher mortality rates observed in *G. mellonella* infected with this strain in comparison with KLEB-33, which demonstrated significantly lower mortality rates despite being a convergent strain (22). However, this model proved inadequate for differentiating between classic and hypervirulent *K. pneumoniae* strains, and thus may not be a reliable indicator of virulence in murine models or in humans in certain cases (48). Additionally, a recent report showed unexpectedly lower virulence in convergent *K. pneumoniae* strains (49). The notable reduction in virulence observed in KLEB-33 *ΔsdiA* supplemented with C6-HSL is consistent with the lower biofilm formation observed in RBB biofilms. It is therefore proposed that C6-HSL may control the expression of virulence and biofilm growth genes independently and in opposition to SdiA, following a hierarchical regulation. This would explain why the biofilm and virulence repressive effect of C6-HSL could only be observed when SdiA was absent.

**Figure 8.**
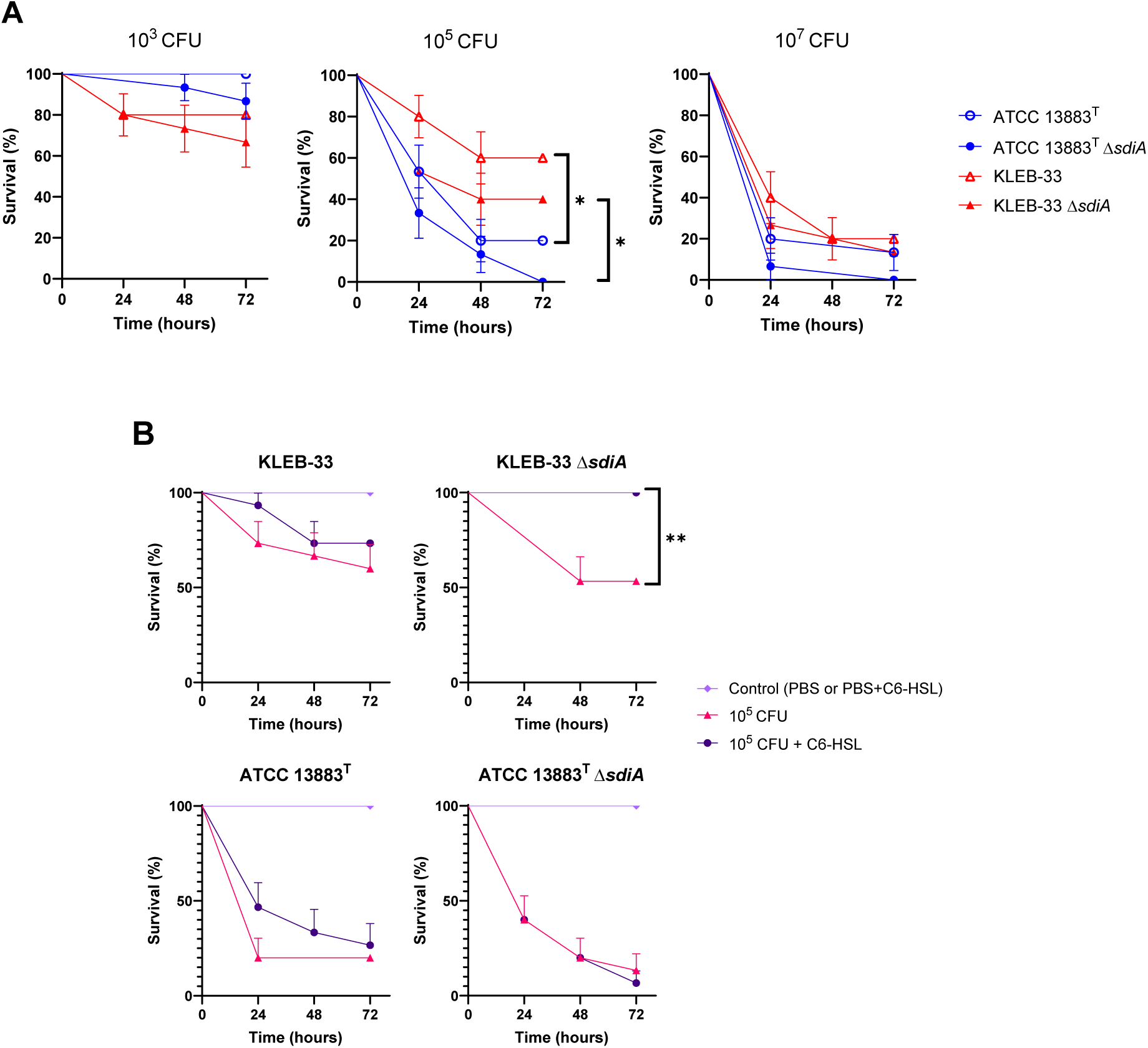
Survival analysis of *Galleria mellonella* following infection with *K. pneumoniae* KLEB-33 and ATCC 13883^T^ wild-type and Δ*sdiA* strains. The survival of the insects was recorded at 24-hour intervals up to 72 hours. The results of the comparison between the concentrations of the strains (**A**) and the effect of C6-HSL addition (5 µM) (**B**) are presented.

## 6 Conclusions

The aim of this study was to elucidate the function of SdiA in the virulence and biofilm formation of *K. pneumoniae*, which is postulated to be the receptor of AHLs. To this end, a series of phenotypes were characterised in SdiA-lacking strains and after the addition of the C6-HSL, which was identified as a biofilm-promoting factor in this bacterium. Our findings appear to indicate that SdiA and C6-HSL influence several traits, with some exhibiting a joint effect, but the majority displaying independent regulation.

The results revealed that SdiA plays a role in repressing biofilm formation, and that its absence was linked to a reduction in resistance to human serum and phage infection, as well as a notable promotion of cell filamentation. However, no impact was observed on macroscopic capsule synthesis. Conversely, the exogenous addition of C6-HSL was found to promote capsule production in a SdiA-dependent manner in one of the strains studied. Moreover, C6-HSL was observed to enhance serum resistance independently of SdiA. Nevertheless, no impact on phage sensitivity was noted. Regarding biofilm formation, C6-HSL has a promoting effect when SdiA is present, but is decreased in its absence. This observation is consistent with the virulence data recorded in *G. mellonella*, which is reduced following the addition of C6-HSL in the absence of SdiA. Additionally, neither SdiA nor C6-HSL affects the composition of the biofilm matrix.

In view of these findings, it seems reasonable to conclude that C6-HSL is not the primary ligand of SdiA in the *K. pneumoniae* strains under consideration. This is because SdiA and C6-HSL are involved in different pathways in some cases, and their effects may even be counteracted by hierarchical regulation. This is corroborated by the observation that certain effects of C6-HSL could only be observed in the absence of SdiA. The observed similarities in the effects of SdiA and C6-HSL on certain phenotypes could lead to the conclusion that they act together, with SdiA acting as a receptor for AHL. However, it is also possible that they act independently. It is important to note that some of the results were dependent on the strain of *K. pneumoniae* studied, as not all the phenotypes observed were consistent in both strains. Our study provides new insights in QS regulation in this pathogen, even so more experiments are necessary to continue with the characterization of their respective ways of action in search of therapeutic targets for the development of new antimicrobial strategies against this species of outstanding clinical interest.

## 9 Supplementary Material Section

**Table S1.**
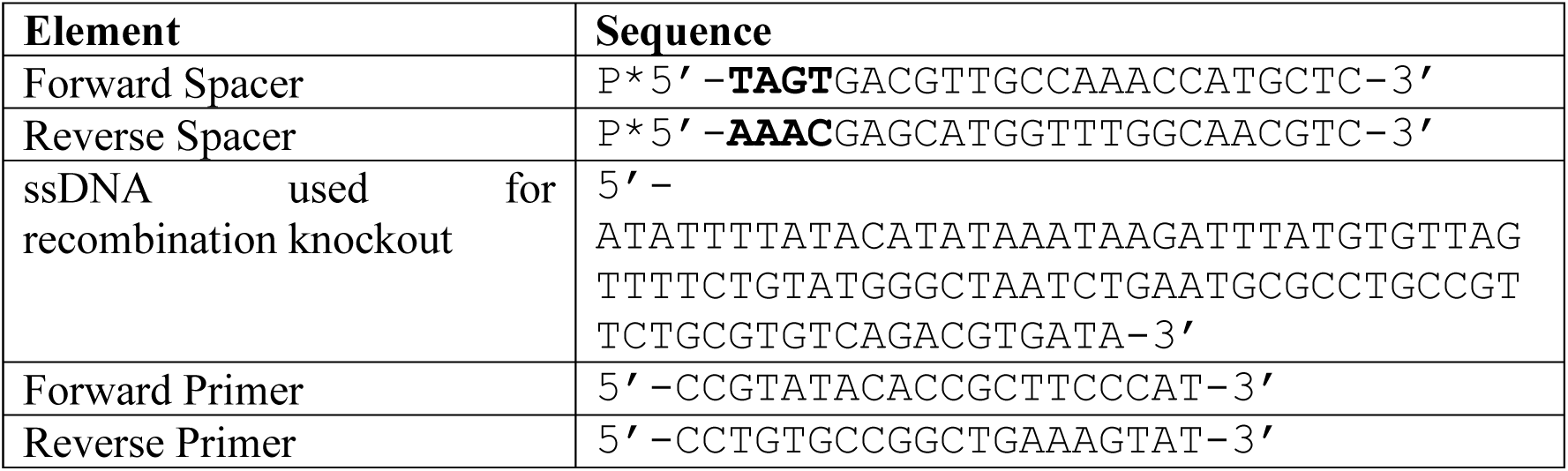
The oligonucleotides employed in this study for the generation of *sdiA* mutants in 1. *K. pneumoniae* are listed below.

**Table S2.** Summary of the QQ activity results recorded from culture samples (supernatant or pellet portions) of *Klebsiella pneumoniae* KLEB-33 and ATCC 13883^T^ wild-type and Δ*sdiA* strains against C6-HSL (10 µM). The remaining AHL signalling activity was evaluated at 6, 12 and 24 hours of incubation. The samples were exposed to biosensor *C. subtsugae* CV026, and the presence or absence of QQ activity was recorded. This entailed observing whether the biosensor exhibited the absence of violacein production, indicative of QQ activity (+), or the presence of violacein production, indicative of negative QQ activity (-). The experiment was repeated twice.

|  | Strain | Condition | Supernatant |  |  |  | Pellet |  |  |  |
| --- | --- | --- | --- | --- | --- | --- | --- | --- | --- | --- |
|  |  |  | 6 h | 12 h | 24 h | 48 h | 6 h | 12 h | 24 h | 48 h |
| *C. subtsugae* CV026 | ATCC 13883<sup>T</sup> | shaking | + | + | + | + | - | - | - | - |
|  |  | static | - | + | + | + | - | - | - | - |
| | ATCC 13883<sup>T</sup> $\Delta sdiA$ | shaking | + | + | + | + | - | - | - | - |
|  |  | static | - | - | + | + | - | - | - | - |
|  | KLEB-33 | shaking | + | + | + | + | - | - | - | - |
|  |  | static | - | + | + | + | - | - | - | - |
| | KLEB-33 $\Delta sdiA$ | shaking | + | + | + | + | - | - | - | - |
|  |  | static | - | + | + | + | - | - | - | - |

**Figure S1.**
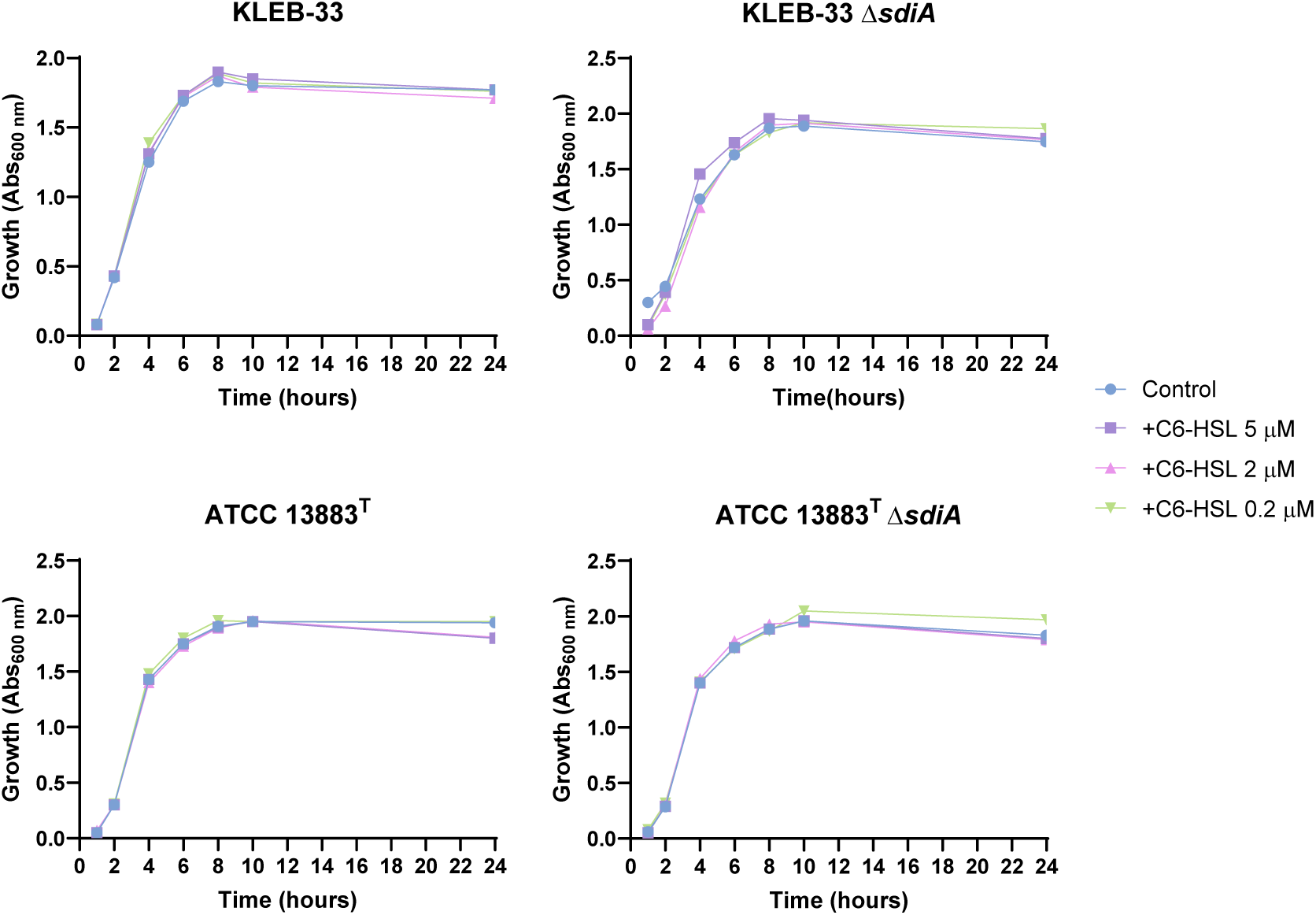
Growth curves of *K. pneumoniae* KLEB-33 and ATCC 13883^T^ wild-type and Δ*sdiA* strains. The impact of C6-HSL supplementation was examined at concentrations of 5, 2, and 0.2 µM.

**Figure S2.**
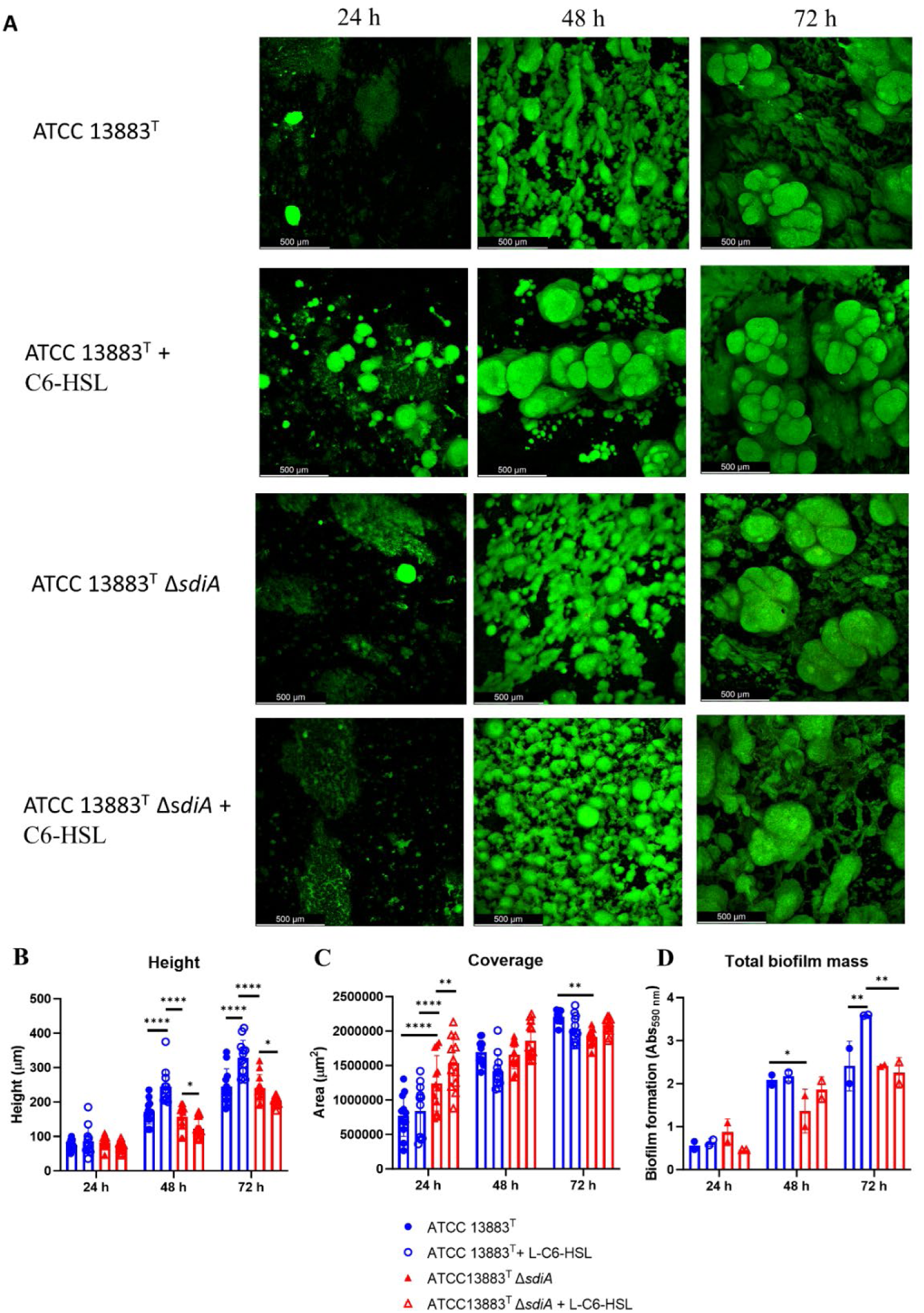
Impact of *sdiA* mutation and AHL addition on biofilm formation in *K. pneumoniae* ATCC 13883^T^. (**A**) Representative CLSM images of biofilms obtained after 24, 48 and 72 h of incubation in the RBB cultivation system and staining with Syto9 fluorescent dye. (**B**) and (**C**) Quantification of height and biofilm coverage from confocal images using ImageJ (v1.54) image analysis software. (**D**) Crystal Violet quantification of wild-type and Δ*sdiA K. pneumoniae* ATCC 13883^T^ strains biofilm biomass after treatment with C6-HSL (5 µM).

**Figure S3.**
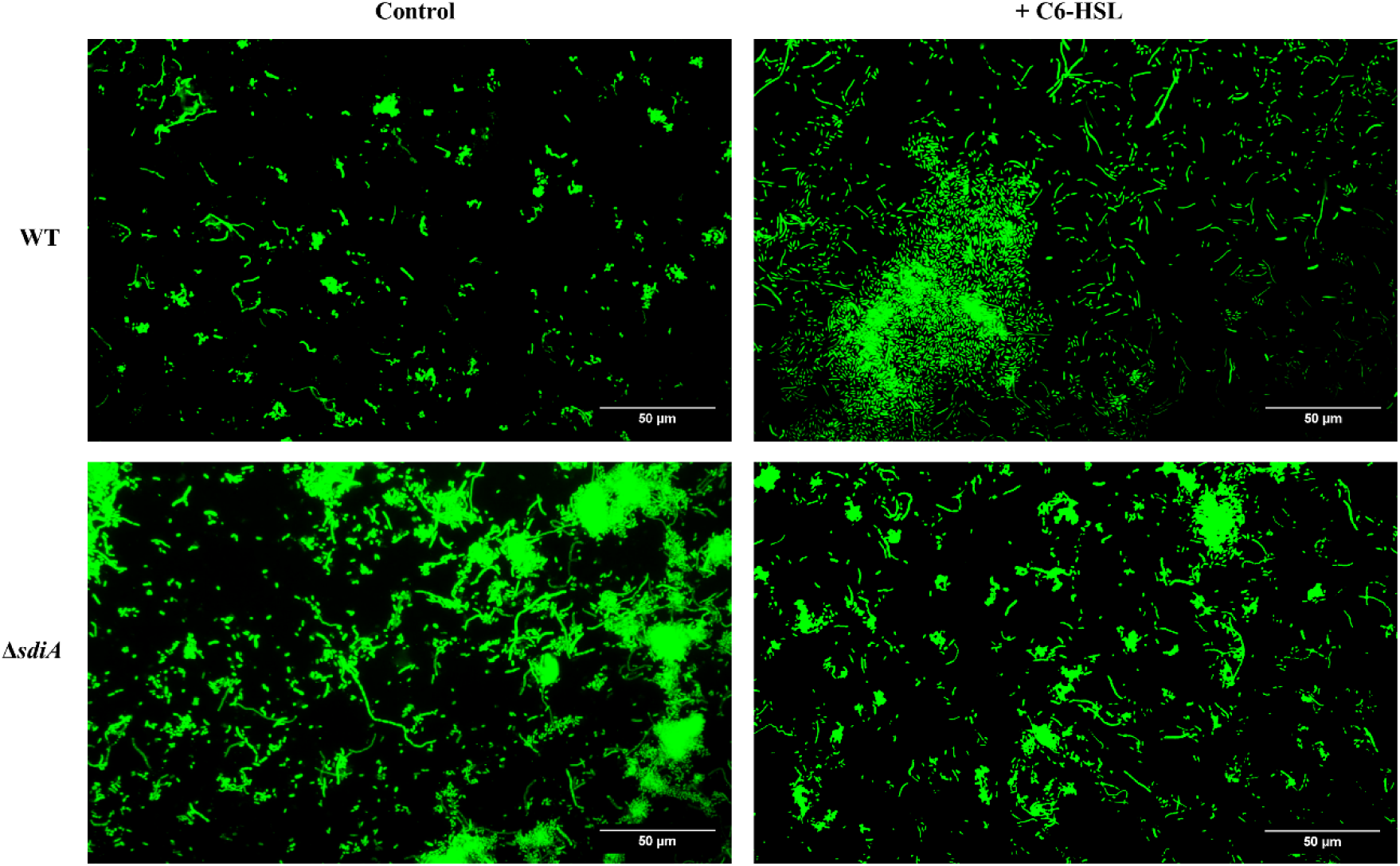
Images illustrating the presence of filamentous bacteria in the initial stages (24 h) of biofilm formation by *K. pneumoniae* KLEB-33. The biofilms were cultivated using the RBB system with or without C6-HSL (5 µM) treatment and subsequently stained with Syto9.

**Figure S4.**
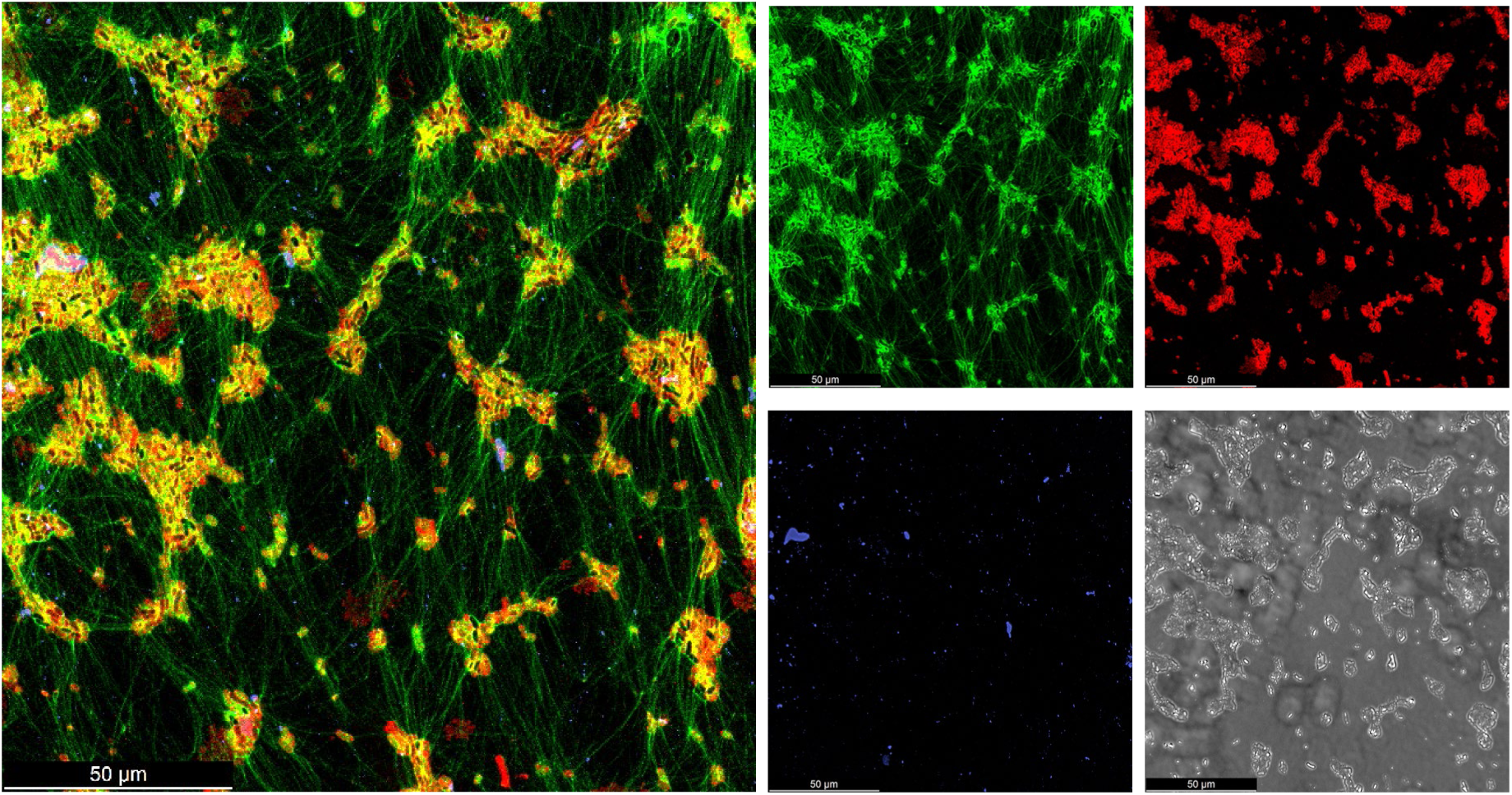
A representative CLSM image of the interior of mushroom-shaped structures observed in *K. pneumoniae* KLEB-33 RBB biofilms indicates that extracellular DNA (eDNA) is a primary component of these biofilms. The image pertains to a 48-hour Δ*sdiA* biofilm. The biofilms were stained with YOYO-1 (eDNA, green), FM-464 (cell membranes, red), and Concanavalin A conjugated with AlexaFluor 594 (exopolysaccharide, blue).

**Figure S5.**
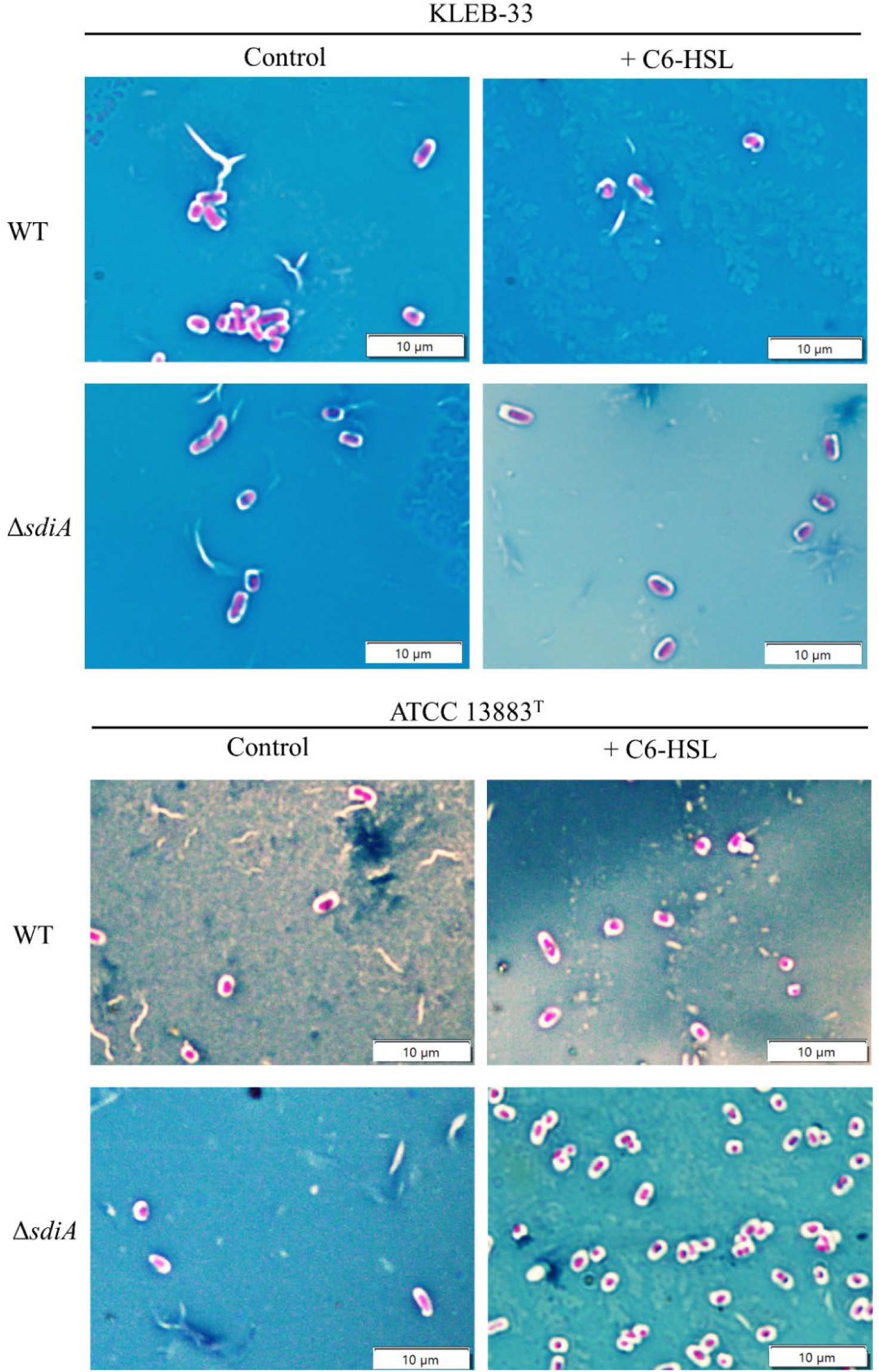
The images displayed represent the results of staining procedures conducted on *Klebsiella pneumoniae* KLEB-33 and ATCC 13883^T^ using the Maneval capsule staining method. The strains under consideration are wild-type (WT) and *ΔsdiA*, with and without the addition of C6-HSL (5 µM). The capsule is revealed as a clear halo between the coloured background (blue) and the stained cell (red).

## References

1. Tacconelli E, Carrara E, Savoldi A, Harbarth S, Mendelson M, Monnet DL, et al. Discovery, research, and development of new antibiotics: the WHO priority list of antibiotic-resistant bacteria and tuberculosis. Lancet Infect Dis. 2018 Mar;18(3):318–27.

4. González-Ferrer S, Peñaloza HF, Budnick JA, Bain WG, Nordstrom HR, Lee JS, et al. Finding Order in the Chaos: Outstanding Questions in Klebsiella pneumoniae Pathogenesis. Ottemann KM, editor. Infect Immun. 2021 Mar 17;89(4):e00693–20.

5. WHO. WHO Bacterial Priority Pathogens List 2024 Bacterial Pathogens of Public Health Importance, to Guide Research, Development, and Strategies to Prevent and Control Antimicrobial Resistance. Geneva: World Health Organization; 2024.

4. Wang S, Payne GF, Bentley WE. Quorum Sensing Communication: Molecularly Connecting Cells, Their Neighbors, and Even Devices. Annu Rev Chem Biomol Eng. 2020 Jun 8;11(1):447–68.

5. Papenfort K, Bassler BL. Quorum sensing signal–response systems in Gram-negative bacteria. Nat Rev Microbiol. 2016 Sep;14(9):576–88.

6. Mayer C, Borges A, Flament-Simon SC, Simões M. Quorum sensing architecture network in *Escherichia coli* virulence and pathogenesis. FEMS Microbiol Rev. 2023 Jul 5;47(4):fuad031.

7. Sabag-Daigle A, Soares JA, Smith JN, Elmasry ME, Ahmer BMM. The Acyl Homoserine Lactone Receptor, SdiA, of *Escherichia coli* and *Salmonella enterica* Serovar Typhimurium Does Not Respond to Indole. Appl Environ Microbiol. 2012 Aug;78(15):5424–31.

8. Rahmati S, Yang S, Davidson AL, Zechiedrich EL. Control of the AcrAB multidrug efflux pump by quorum-sensing regulator SdiA. Mol Microbiol. 2002 Feb;43(3):677–85.

9. Suzuki K, Wang X, Weilbacher T, Pernestig AK, Melefors Ö, Georgellis D, et al. Regulatory Circuitry of the CsrA/CsrB and BarA/UvrY Systems of *Escherichia coli*. J Bacteriol. 2002 Sep 15;184(18):5130–40.

10. Król JE, Hall DC, Balashov S, Pastor S, Sibert J, McCaffrey J, et al. Genome rearrangements induce biofilm formation in *Escherichia coli* C – an old model organism with a new application in biofilm research. BMC Genomics. 2019 Dec;20(1):767.

11. Cao Y, Li L, Zhang Y, Liu F, Xiao X, Li X, et al. Evaluation of *Cronobacter sakazakii* biofilm formation after sdiA knockout in different osmotic pressure conditions. Food Res Int. 2022 Jan;151:110886.

12. Lindsay A, Ahmer BMM. Effect of *sdiA* on Biosensors of *N* -Acylhomoserine Lactones. J Bacteriol. 2005 Jul;187(14):5054–8.

13. Lee J, Jayaraman A, Wood TK. Indole is an inter-species biofilm signal mediated by SdiA. BMC Microbiol. 2007 Dec;7(1):42.

14. Ghosh D, Roy K, Williamson KE, Srinivasiah S, Wommack KE, Radosevich M. Acyl-Homoserine Lactones Can Induce Virus Production in Lysogenic Bacteria: an Alternative Paradigm for Prophage Induction. Appl Environ Microbiol. 2009 Nov 15;75(22):7142–52.

15. Janssens JCA, Metzger K, Daniels R, Ptacek D, Verhoeven T, Habel LW, et al. Synthesis of *N* -Acyl Homoserine Lactone Analogues Reveals Strong Activators of SdiA, the *Salmonella enterica* Serovar Typhimurium LuxR Homologue. Appl Environ Microbiol. 2007 Jan 15;73(2):535–44.

16. Styles MJ, Early SA, Tucholski T, West KHJ, Ge Y, Blackwell HE. Chemical Control of Quorum Sensing in *E. coli* : Identification of Small Molecule Modulators of SdiA and Mechanistic Characterization of a Covalent Inhibitor. ACS Infect Dis. 2020 Dec 11;6(12):3092–103.

17. Pacheco T, Gomes AÉI, Siqueira NMG, Assoni L, Darrieux M, Venter H, et al. SdiA, a Quorum-Sensing Regulator, Suppresses Fimbriae Expression, Biofilm Formation, and Quorum-Sensing Signaling Molecules Production in *Klebsiella pneumoniae*. Front Microbiol. 2021 Jun 21;12:597735.

18. Subramoni S, Venturi V. LuxR-family ‘solos’: bachelor sensors/regulators of signalling molecules. Microbiology. 2009 May 1;155(5):1377–85.

19. Hosny RA, Fadel MA. Detection of Quorum Sensing N-Acyl-Homoserine Lactone Molecules Produced by Different Resistant *Klebsiella pneumoniae* Isolates Recovered from Poultry and Different Environmental Niches. Appl Biochem Biotechnol. 2021 Oct;193(10):3351–70.

20. Ngeow Y, Cheng H, Chen J, Yin WF, Chan KG. Short Chain N-Acylhomoserine Lactone Production by Clinical Multidrug Resistant *Klebsiella pneumoniae* Strain CSG20. Sensors. 2013 Nov 7;13(11):15242–51.

21. Wang H, Cai T, Weng M, Zhou J, Cao H, Zhong Z, et al. Conditional production of acyl-homoserine lactone-type quorum-sensing signals in clinical isolates of enterobacteria. J Med Microbiol. 2006 Dec 1;55(12):1751–3.

22. Silva-Bea S, García-Meniño I, Rey S, Romero M, Fernández J, Hammerl JA, et al. Draft genome sequence of Klebsiella pneumoniae KLEB-33: a convergent biofilm hyperforming multiresistant strain belonging to the emerging ST16 lineage harboring multiple hypervirulence genes. Putonti C, editor. Microbiol Resour Announc. 2024 Apr 11;13(4):e01192–23.

23. Wang Y, Wang S, Chen W, Song L, Zhang Y, Shen Z, et al. CRISPR-Cas9 and CRISPR-Assisted Cytidine Deaminase Enable Precise and Efficient Genome Editing in Klebsiella pneumoniae. Drake HL, editor. Appl Environ Microbiol. 2018 Dec;84(23):e01834–18.

24. Miles AA, Misra SS, Irwin JO. The estimation of the bactericidal power of the blood. Epidemiol Infect. 1938 Nov;38(6):732–49.

25. Silva-Bea S, Romero M, Parga A, Fernández J, Mora A, Otero A. Comparative analysis of multidrug-resistant *Klebsiella pneumoniae* strains of food and human origin reveals overlapping populations. Int J Food Microbiol. 2024 Mar;413:110605.

26. Romero M, Mayer C, Heeb S, Wattanavaekin K, Cámara M, Otero A, et al. Mushroom-shaped structures formed in *Acinetobacter baumannii* biofilms grown in a roller bioreactor are associated with quorum sensing–dependent Csu-pilus assembly. Environ Microbiol. 2022 Sep;24(9):4329–39.

27. Exterkate RAM, Crielaard W, Ten Cate JM. Different Response to Amine Fluoride by *Streptococcus mutans* and Polymicrobial Biofilms in a Novel High-Throughput Active Attachment Model. Caries Res. 2010;44(4):372–9.

28. Parga A, Muras A, Otero-Casal P, Arredondo A, Soler-Ollé A, Àlvarez G, et al. The quorum quenching enzyme Aii20J modifies in vitro periodontal biofilm formation. Front Cell Infect Microbiol. 2023 Feb 2;13:1118630.

29. Dorman MJ, Feltwell T, Goulding DA, Parkhill J, Short FL. The Capsule Regulatory Network of Klebsiella pneumoniae Defined by density-TraDISort. Chang YF, editor. mBio. 2018 Dec 21;9(6):e01863–18.

30. Hughes RB, Smith AC. Capsule-Stain-Protocols [Internet]. American Society for Microbiology; 2007. Available from: https://asm.org/ASM/media/Protocol-Images/Capsule-Stain-Protocols.pdf?ext=.pdf

31. Lv J, Zhu J, Wang T, Xie X, Wang T, Zhu Z, et al. The Role of the Two-Component QseBC Signaling System in Biofilm Formation and Virulence of Hypervirulent *Klebsiella pneumoniae* ATCC43816. Front Microbiol. 2022 Apr 6;13:817494.

32. Xie Y, Wahab L, Gill J. Development and Validation of a Microtiter Plate-Based Assay for Determination of Bacteriophage Host Range and Virulence. Viruses. 2018 Apr 12;10(4):189.

33. Gato E, Vázquez-Ucha JC, Rumbo-Feal S, Álvarez-Fraga L, Vallejo JA, Martínez-Guitián M, et al. Kpi, a chaperone-usher pili system associated with the worldwide-disseminated high-risk clone *Klebsiella pneumoniae* ST-15. Proc Natl Acad Sci. 2020 Jul 21;117(29):17249–59.

34. Insua JL, Llobet E, Moranta D, Pérez-Gutiérrez C, Tomás A, Garmendia J, et al. Modeling Klebsiella pneumoniae Pathogenesis by Infection of the Wax Moth Galleria mellonella. Bliska JB, editor. Infect Immun. 2013 Oct;81(10):3552–65.

35. Panchal J, Prajapati J, Dabhi M, Patel A, Patel S, Rawal R, et al. Comprehensive computational investigation for ligand recognition and binding dynamics of SdiA: a degenerate LuxR -type receptor in Klebsiella pneumoniae. Mol Divers [Internet]. 2024 Jan 12 [cited 2024 Feb 10]; Available from: https://link.springer.com/10.1007/s11030-023-10785-6

36. Milton DL, Chalker VJ, Kirke D, Hardman A, Cámara M, Williams P. The LuxM Homologue VanM from *Vibrio anguillarum* Directs the Synthesis of *N* -(3-Hydroxyhexanoyl)homoserine Lactone and *N* -Hexanoylhomoserine Lactone. J Bacteriol. 2001 Jun 15;183(12):3537–47.

37. Gopu V, Meena CK, Murali A, Shetty PH. Petunidin as a competitive inhibitor of acylated homoserine lactones in *Klebsiella pneumoniae*. RSC Adv. 2016;6(4):2592–601.

38. Chan KG. Expression of *Klebsiella* sp. lactonase *ahlK* gene is growth-phase, cell-population density and *N* -acylhomoserine lactone independent. Front Life Sci. 2013 Dec;7(3–4):132–9.

39. Abell-King C, Costas A, Duggin IG, Söderström B. Bacterial filamentation during urinary tract infections. Coers J, editor. PLOS Pathog. 2022 Dec 1;18(12):e1010950.

40. Das T, Manefield M. Pyocyanin Promotes Extracellular DNA Release in Pseudomonas aeruginosa. Rohde H, editor. PLoS ONE. 2012 Oct 8;7(10):e46718.

41. Yang N, Lan L. *Pseudomonas aeruginosa* Lon and ClpXP proteases: roles in linking carbon catabolite repression system with quorum-sensing system. Curr Genet. 2016 Feb;62(1):1–6.

42. Campoccia D, Montanaro L, Arciola CR. Extracellular DNA (eDNA). A Major Ubiquitous Element of the Bacterial Biofilm Architecture. Int J Mol Sci. 2021 Aug 23;22(16):9100.

43. Mann EE, Rice KC, Boles BR, Endres JL, Ranjit D, Chandramohan L, et al. Modulation of eDNA Release and Degradation Affects Staphylococcus aureus Biofilm Maturation. Ratner AJ, editor. PLoS ONE. 2009 Jun 9;4(6):e5822.

44. Nunez C, Kostoulias X, Peleg AY, Short F, Qu Y. A comprehensive comparison of biofilm formation and capsule production for bacterial survival on hospital surfaces. Biofilm. 2023 Dec;5:100105.

45. Majkowska-Skrobek G, Markwitz P, Sosnowska E, Lood C, Lavigne R, Drulis-Kawa Z. The evolutionary trade-offs in phage-resistant *Klebsiella pneumoniae* entail cross-phage sensitization and loss of multidrug resistance. Environ Microbiol. 2021 Dec;23(12):7723–40.

46. Tang M, Huang Z, Zhang X, Kong J, Zhou B, Han Y, et al. Phage resistance formation and fitness costs of hypervirulent *Klebsiella pneumoniae* mediated by K2 capsule-specific phage and the corresponding mechanisms. Front Microbiol. 2023 Jul 19;14:1156292.

47. Huja S, Oren Y, Biran D, Meyer S, Dobrindt U, Bernhard J, et al. Fur Is the Master Regulator of the Extraintestinal Pathogenic Escherichia coli Response to Serum. Rappuoli R, editor. mBio. 2014 Aug 29;5(4):e01460–14.

48. Russo TA, MacDonald U. The Galleria mellonella Infection Model Does Not Accurately Differentiate between Hypervirulent and Classical Klebsiella pneumoniae. Papasian CJ, editor. mSphere. 2020 Feb 26;5(1):e00850–19.

49. Kochan TJ, Nozick SH, Valdes A, Mitra SD, Cheung BH, Lebrun-Corbin M, et al. *Klebsiella pneumoniae* clinical isolates with features of both multidrug-resistance and hypervirulence have unexpectedly low virulence. Nat Commun. 2023 Dec 2;14(1):7962.

